# rDNA copy number variation in yeast alters response to environmental conditions

**DOI:** 10.1101/2024.09.30.615902

**Authors:** Kevin Thornton, Elizabeth X. Kwan, Kerry Bubb, Luana Paleologu, M. K. Raghuraman, Bonita J. Brewer, Josh T. Cuperus, Christine Queitsch

## Abstract

Ribosomal DNA (rDNA) in eukaryotes is maintained in hundreds of copies with rDNA copy number varying greatly among individuals within a species. In the budding yeast *Saccharomyces cerevisiae*, the rDNA copy number across wild isolates ranges from 90 to 300 copies. Previous studies showed that 35 rDNA copies are sufficient for ribosome biogenesis in this yeast and enable wild-type-like growth in standard laboratory growth conditions. We addressed two major questions concerning rDNA copy number variation in this yeast: (1) What are the fitness consequences of rDNA copy number variation outside and within the natural range in standard laboratory growth conditions? (2) Do these fitness effects change in different growth conditions? We used growth competitions to compare the fitness effects of rDNA copy number variation in otherwise isogenic strains whose rDNA copy number ranged from 35 to 200. In standard growth conditions, we found that fitness gradually increases from 35 rDNA copies until reaching a plateau that spans from 98 to 160 rDNA copies, well within the natural range. However, rDNA copy number-dependent fitness differed across environments. Compared to standard growth conditions, strains with higher rDNA copy numbers showed improved fitness when grown at increased temperature or in glycerol media. Moreover, the gradual fitness increase with increasing rDNA copy number in standard growth conditions gave way to a markedly lower fitness of strains with copy numbers below the natural range in these two stress conditions. These results suggest that selective pressures drive rDNA copy number in this yeast to at least ~100 copies and that a higher number of copies buffers against environmental stress. The similarity of the *S. cerevisiae* rDNA copy number range to the ranges reported in *C. elegans, D. melanogaster*, and humans points to conserved selective pressures maintaining the range of natural rDNA copy number in these highly diverse species.

## INTRODUCTION

Ribosomal DNA (rDNA) encodes the ribosomal RNAs that provide the structural and enzymatic features of ribosomes. The rDNA is maintained in tandem arrays at hundreds of copies per cell (Kobayashi *et al*. 1998; Nelson *et al*. 2019). The repetitive nature of the rDNA arrays makes them prone to copy number variation within species. There are 90 to 300 copies in the yeast *Saccharomyces cerevisiae*, 70 to 400 copies in the worm *Caenorhabditis elegans*, 80 to 600 copies in the fly *Drosophila melanogaster*, 500 to 2,500 copies in the plant *Arabidopsis thaliana*, and 100 to 600 copies in humans (Mohan and Ritossa 1970; Thompson *et al*. 2013; Morton *et al*. 2020, 2023; Hall *et al*. 2021, 2022). Given the breadth of variation, both in copy number and the fraction of the genome it represents, rDNA may be an underappreciated source of genetic variation influencing fitness.

rDNA copy number variation is increasingly recognized as an influential factor in cell physiology. A minimum number of rDNA copies is required for the high level of rRNA transcription necessary for ribosome biogenesis with rRNAs making up approximately 80% of total cellular RNA in yeast (Warner 1999). For *S. cerevisiae*, this minimum number of rDNA copies is 35 (French *et al*. 2003; Kim *et al*. 2006; Ide *et al*. 2010; Kwan *et al*. 2023), a number that is significantly lower than observed in wild yeast isolates and laboratory strains (Morton *et al*. 2020; Hall *et al*. 2022). Strains with substantially reduced rDNA copy numbers sufficient for ribosome biogenesis (35-40 copies) show compromised genome replication and increased susceptibility to DNA damage but no growth defects in standard laboratory growth conditions (Ide *et al*. 2010; Salim *et al*. 2017; Kwan *et al*. 2023). As few as 8-10 rDNA copies suffice to maintain yeast viability (Sanchez *et al*. 2019; Jiang *et al*. 2024); however the resulting limitation on ribosome production creates a strong selective pressure for suppressors that ameliorate their slow-growth phenotype (Sanchez et al., 2019) In animals and humans, reductions of rDNA copy number below the naturally occurring range of variation is associated with developmental abnormalities and disease phenotypes (Ritossa and Atwood 1966; Xu *et al*. 2017; Valori *et al*. 2020; Morton *et al*. 2023). However, the intrinsic differences among tissues in multicellular animals makes it difficult to determine the extent to which rDNA copy number variation affects cellular fitness.

We therefore wanted to investigate in yeast cells (1) if rDNA copy number variation below and within the natural range affects cellular fitness; (2) if increases in rDNA copy number from the minimal copy number up to levels observed in wild and laboratory strains affect fitness gradually or if there are copy number thresholds; and (3) what additional biological processes are affected by substantial reduction in rDNA copy number. To tackle these questions, we generated a set of isogenic strains in the yeast *S. cerevisiae* containing between 35 to 200 copies of rDNA, which spans the range from the threshold required for ribosome biogenesis up to the natural variation present in the species. We examined these strains by competitive fitness assays in different growth conditions, and used transcriptome analysis of a strain with 35 rDNA copies and a control strain with 180 copies to identify additional, potentially altered pathways.

We found that rDNA copy number variation profoundly affects fitness in haploid yeast. In standard laboratory conditions, strains below ~100 copies showed reduced fitness that correlated with the severity of copy number reduction. In contrast, strains with copy numbers ranging from 98 to 160 rDNA copies displayed similarly high fitness, establishing a fitness plateau that roughly coincides with the naturally occurring rDNA copy number range reported for this yeast (Morton *et al*. 2020; Hall *et al*. 2022). Additional rDNA copies are detrimental, as fitness decreased in strains with more than 160 rDNA copies. A comparison of the transcriptomes of the 35 and 180 rDNA copy number strains showed the expected altered expression of genes involved in mitigating DNA replication stress, in addition to unexpected expression changes in canonical stress response genes. We followed up on the latter findings with phenotyping and competition assays at increased temperature and with growth on a nonfermentable carbon source. We found that the rDNA copy number variants showed profoundly different fitness in different growth conditions, and that the gradual fitness changes observed in standard growth conditions had given way to threshold effects. Taken together, our findings suggest that the range of rDNA copy numbers observed in wild yeast strains acts as a buffer under varied environments.

## RESULTS

### rDNA copy number affects yeast fitness in standard growth conditions

To generate a set of isogenic *S. cerevisiae* strains with variable rDNA copy numbers, we started with a *fob1Δ* strain that stably maintains 35 copies in a S288c strain background (Kobayashi *et al*. 1998; Ide *et al*. 2010; Kwan *et al*. 2023). After a transient re-introduction of *FOB1* to allow for copy number expansion, we obtained multiple clones for strains with copy numbers ranging from 45 to 180. A strain with 200 rDNA copies was isolated by chance during a cross. We verified rDNA copy number for three clones of each strain by CHEF gel electrophoresis and Southern blotting (**Figure 1A,B**; Tsuchiyama *et al*. 2013; Kwan *et al*. 2016; Morton *et al*. 2020).

**Figure 1:**
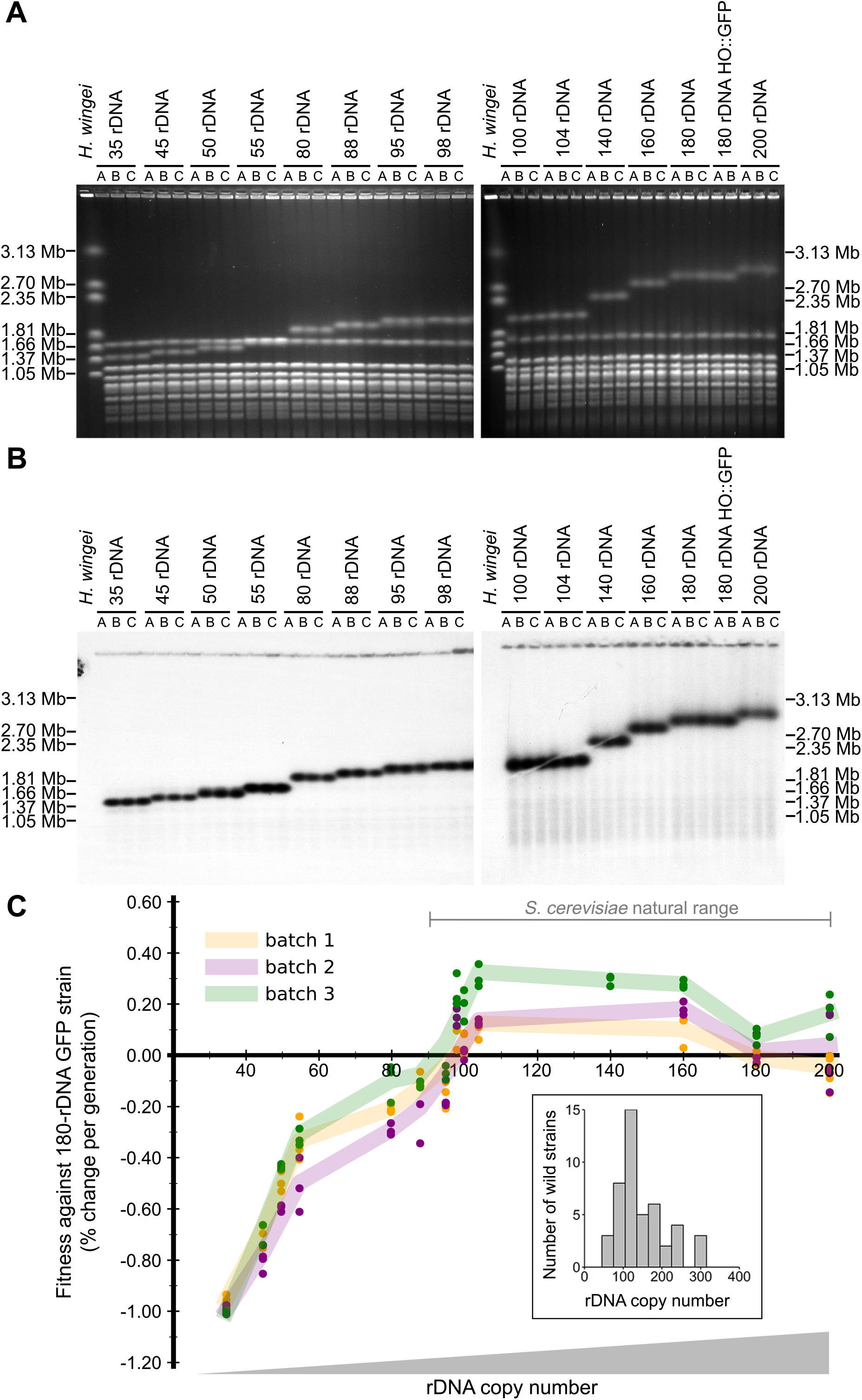
rDNA copy number affects fitness in standard growth conditions. CHEF gel electrophoresis was performed to verify rDNA copy number in the triplicate clones used to inoculate fitness competitions under standard growth conditions. (A) Ethidium bromide-stained gel and (B) the resulting Southern blot hybridized with a single copy Chr. XII probe (*CDC45*). (C) Graph of fitness values calculated by the rate of population change for each rDNA copy number test strain against the GFP competitor strain (180 rDNA copies). Dots indicate individual fitness calculated of three replicates for each strain; shared dot and line colors indicate each of three experimental replicates. Inset: distribution of rDNA copy genotypes in wild *S. cerevisiae* isolates.

Strains with copy numbers between 35 and 200 showed similar growth rates (**Figure S1A**), consistent with previous observations (Jiang *et al*. 2024). To magnify small phenotypic differences between strains (Conti *et al*. 2022), we performed competition assays to assess the relative fitness of strains with varying rDNA copy numbers. The competitor strain for each experiment was a GFP-tagged strain with 180 rDNA copies that was otherwise isogenic. To determine the competitive fitness of each strain against the GFP 180 rDNA copy number strain, we mixed the two starting cultures in triplicate in a 1:1 ratio and grew them to saturation (Gresham and Dunham 2014). Every 24 hours, we took samples to measure the ratio of GFP-positive to GFP-negative cells by flow cytometry. We then diluted the competition cultures 1:1000 into fresh media and grew them again to saturation, allowing for approximately 10 generations of growth. We repeated this process for ~50 generations, and estimated fitness by tracking the ratios of the GFP competitor strain to each test strain over the indicated number of cell divisions. The comparison of the competitor GFP 180 rDNA copy number strain with the 180 rDNA copy number strain yielded an estimate for the fitness cost of GFP expression of 0.08% per generation (**Figure 1C**).

We noticed three trends in the fitness data: (1) reduced fitness in strains with fewer than 98 rDNA copies; (2) a fitness plateau for strains with 98 to 160 rDNA copies; and (3) subtly reduced fitness in strains with more than 160 rDNA copies. The strain with the fewest rDNA copies (35 copies) showed the greatest fitness defect (**Figure 1C**). Strain fitness appeared to increase in a tri-phasic pattern with steep gains as rDNA copies increased from 35 to 55, more gradual fitness gains from 55 to 88 copies, and steep fitness gains from 88 to 104 copies. There was a sharp fitness threshold around 100 rDNA copies, followed by an extended fitness plateau. The fitness plateau, encompassing strains with 98 to 160 rDNA copies (**Figure 1C**), mirrored the distribution of rDNA copy number genotypes in wild *S. cerevisiae* isolates (**Figure 1C, inset**). Additional rDNA copies above 160 resulted in moderately reduced fitness. A strain with 180 rDNA copies showed reduced fitness compared to the strains with 98 to 160 rDNA copies, but greater fitness than the strains with fewer than 98 rDNA copies (**Figure 1C**). Strains with 200 rDNA copies also showed moderately reduced fitness compared to the strains with 98 to 160 rDNA copies (**Figure 1C**). We conclude that rDNA copy number variants with fewer than 98 and more than 160 rDNA copies in the S288c strain background show reduced fitness under standard growth conditions. Strain fitness did not increase linearly but showed strong threshold effects, coinciding with the lower boundary of the natural variation range across *S. cerevisiae* strains.

### rDNA expansion upon FOB1 re-introduction into a short rDNA strain slows upon reaching ~100 rDNA copies

Strains with substantially reduced rDNA arrays increase their rDNA copy numbers back to wild-type levels when *FOB1* is re-introduced (Kobayashi *et al*. 1998). The re-introduction of *FOB1* and passaging of cells to an rDNA copy number equilibrium enables another, possibly more nuanced analysis of the fitness effects of rDNA copy number than in the described competition experiments. Instead of competing two strains with fixed rDNA copy numbers, the *FOB1* re-introduction experiment produces naturally occurring variation in rDNA copy number among the cells in the culture and allows for selection among different rDNA copy number variants. Moreover, assessing fitness of wild-type *FOB1* cells avoids any possible effects of the *fob1Δ* mutation. We hypothesized that the expansion of rDNA arrays upon *FOB1* re-introduction would slow when cells reached ~100 copies in standard growth conditions, which would be consistent with fitness being the primary driver for rDNA expansion. Alternatively, rDNA copy number might expand to the 150-180 rDNA copies reported for this strain background (Ide *et al*. 2013; Kwan *et al*. 2016; Morton *et al*. 2020), supporting a model in which strain-specific rDNA copy number is determined by a counting mechanism (Iida and Kobayashi 2019a; b).

To conduct this experiment, we re-introduced *FOB1* by crossing a 35 rDNA *fob1Δ* strain with a 170 rDNA *FOB1* strain, picked three 35 rDNA *FOB1* spores, and passaged the resulting cultures (and the two parental strains) for 300 generations. Samples were collected every 30 generations for subsequent CHEF gel electrophoresis and Southern blotting to determine rDNA copy number (**Figure 2A, B**). After ~30 generations, the three spore-derived cultures showed an average of ~56 rDNA copies (**Figure 2C**). After 150 generations, they had ~100 rDNA copies, and ~107 rDNA copies after 300 generations, suggesting that rDNA copy number expansion slows once ~100 rDNA copies are reached. As expected, the parental strains, 35 rDNA *fob1Δ* and 170 rDNA *FOB1*, did not change significantly in their respective rDNA copy numbers throughout this experiment (**Figure 2C**). We conclude that there is no significant fitness advantage in having more than ~100 rDNA copies in standard growth conditions, consistent with our observation of a sharp fitness increase associated with this approximate copy number.

**Figure 2:**
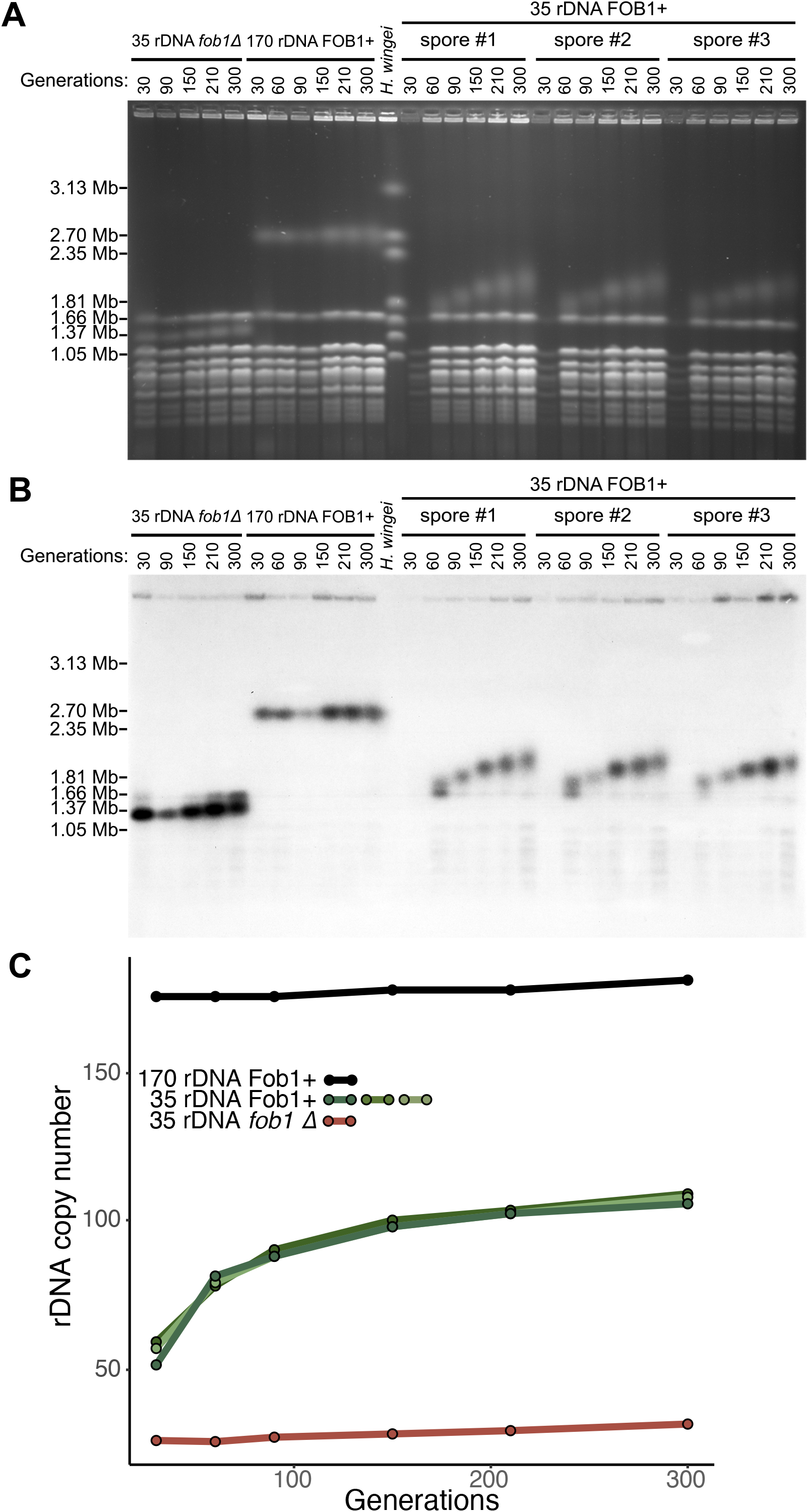
rDNA expansion decelerates upon reaching ~100 rDNA copies. Examination of chromosome XII size by (A) CHEF gel electrophoresis and (B) Southern blotting. (C) rDNA copy numbers from each sample were estimated from the above CHEF gel and plotted against generation number.

### A strain with 35 rDNA copies shows altered gene expression specifically in late S-phase

Strains with substantial reduction of rDNA copy number (20-40 copies) show increased sensitivity to mutagens, defects in genome replication, and cell cycle misregulation (Ide *et al*. 2010; Kwan *et al*. 2023). Although these phenotypic effects might suffice to cause the observed fitness defects, we used RNA-seq analysis to discover other biological processes that might be altered between a short (35 copies) and a control (180 copies) rDNA strain. Because low rDNA copy number causes genome replication delays (Kwan *et al*. 2023), we collected cells in late S-phase (40 minutes after synchronization) and in asynchronous, logarithmically-growing cells. We identified only six genes that were differentially expressed (DEGs) between the 35 and the 180 rDNA strains in asynchronous cultures (p-adj < 0.01), and 708 genes that were differentially expressed between these strains in late S-phase, with three genes overlapping between the two sets (**Figure 3**). Of these three, *PRP11*, implicated in splicing, and *STR3*, implicated in peroxisome function, were expressed at lower levels in the 35 rDNA strain. The third gene, *HSP12*, was expressed at higher levels in the 35 rDNA strain. This gene is induced in response to many stressful conditions, including heat stress, ethanol and salt stress, in addition to DNA replication stress. The small number of genes with expression differences between the 35 and the 180 rDNA strain in asynchronous culture is consistent with previous observations that strains with these rDNA copy numbers do not differ significantly in growth rates, rRNA expression, or ribosome function (Mohan and Ritossa 1970; French *et al*. 2003; Kim *et al*. 2006; Ide *et al*. 2010; Kwan *et al*. 2023).

**Figure 3.**
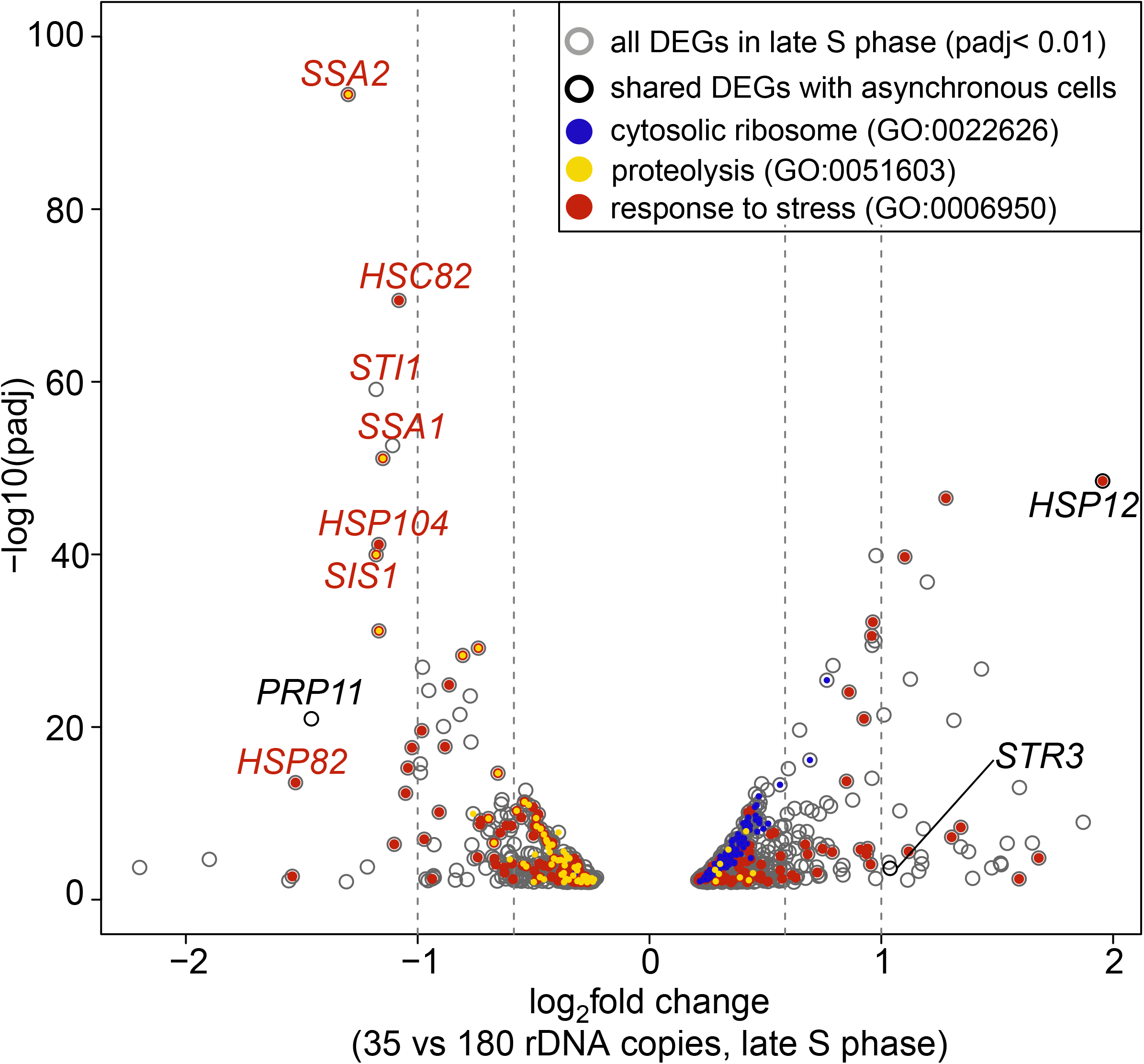
Transcriptome analysis reveals expression changes specific to late S phase. Genes with significantly different expression (padj<0.01) between the 35 rDNA copy number strain and the 180 rDNA copy number strain in late S phase. The three genes that were also differentially expressed in asynchronous growth conditions are represented by black circles; two genes are labeled in black. *HSP12* is labeled in red because of its function in the stress response. Genes within select GO categories are indicated with colored dots: blue for cytosolic ribosome (GO:0022626), yellow for proteolysis involved in protein catabolic process (GO:0051603), and red for response to stress (GO:0006950). Many canonical heat stress proteins and their co-chaperones were significantly downregulated in the 35 rDNA copy number strain in late S phase (labeled in red).

However, in late S-phase, genes upregulated in the 35 rDNA strain were enriched for GO terms related to ribosomal structure and assembly. For example, 115 of the 420 upregulated genes in the 35 rDNA strain belonged to the ‘cytosolic ribosome’ category. In contrast, the 288 genes downregulated in the 35 rDNA copy strain were enriched for GO categories related to protein degradation (*e*.*g*., proteolysis, response to stress, and proteasome complex).

To gain further insight into the significance of these S-phase-specific expression changes, we focused on genes with at least a 1.5-fold change in expression. We found 88 genes upregulated in the 35 rDNA strain (adjpval<1e-6), with GO term enrichments for structural constituent of chromatin, protein heterodimerization activity, and nucleosome. The top ten most upregulated genes were *HSP12*; five small nucleolar RNAs involved in rDNA processing; *PCNA*, a sliding replication clamp; *DDR2*, a multi-stress response gene named for its function in the DNA damage response; *SOE1*, a tRNA gene and suppressor of *CDC8*, a kinase functioning in the biosynthesis of deoxyribonucleotides; and *SIP18*, a gene involved in salt stress.

There were 83 downregulated genes in the 35 rDNA strain (adj-pval<1e-7), with GO term enrichments for protein folding, response to stress, and MCM helicase complex. The latter is consistent with the genome replication defects in the 35 rDNA strain: MCM helicase is essential for DNA replication initiation and elongation and is recruited to the origins of DNA replication as part of the pre-replicative complex (Bell and Labib 2016). The top ten most strongly downregulated genes included both HSP70s (*SSA1* and *SSA2)*; *SIS1*, a co-chaperone of HSP40; *STI1*, a co-chaperone of HSP90; and *HSP104*, a disaggregase functioning with HSP70 and HSP40 required for the acquisition of thermotolerance (Lindquist and Kim 1996). The other genes induced as part of the canonical heat stress response, the two HSP90s (HSC82 and HSP82) were the 13^th^ and 15^th^ most strongly downregulated genes, respectively. While some of the differentially expressed genes and GO annotations are consistent with prior studies (Ide *et al*. 2007, 2010; Salim *et al*. 2017; Kwan *et al*. 2023), the unexpected S-phase-specific finding of altered chaperone expression warranted further investigation as to whether rDNA copy number variation might play a role in response to stress or altered growth conditions.

### rDNA copy number variants show altered fitness at 37°C and in glycerol media

To determine whether rDNA copy number variation might cause phenotypic effects in response to heat stress, we performed competition experiments at 37°C. Results differed markedly from those observed in standard growth conditions. Specifically, the gradual fitness increase for strains with up to 98 copies in standard conditions gave way to a plateau of low fitness for strains with copy numbers ranging from 45 to 95 (**Figure 4A**). Although the 35 rDNA copy number strain showed by far the greatest fitness defect among the rDNA copy number variants, compared to its fitness in standard laboratory conditions, it was little affected by heat stress (**Figure 4A, Figure S2**). Strains with higher copy numbers, from 80 to 95 copies, showed sharply decreased fitness in response to heat stress compared to their performance in standard growth conditions (**Figure 4A**). In contrast, the strains with 180 and 200 rDNA copies showed greater fitness in response to heat stress than in standard growth conditions (**Figure 4A**). To rule out the possibility that heat stress altered rDNA copy numbers during the course of these experiments, we examined CHEF gels and Southern blots of all strains at the start of competitions (day 0) and at the end of competitions (day 5). rDNA copy numbers did not change during the heat stress competitions (**Figure S3**).

**Figure 4:**
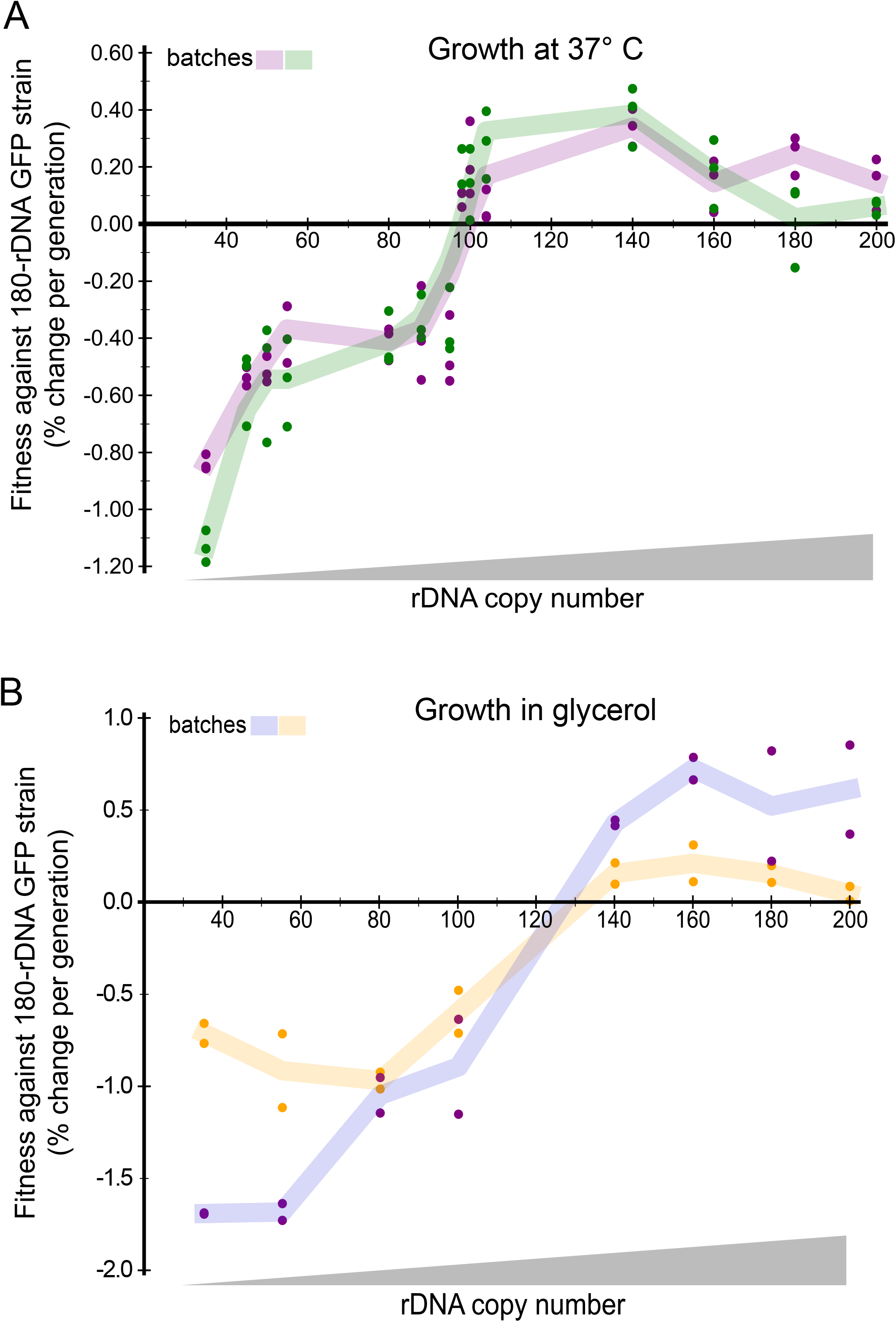
rDNA copy number variants show altered fitness during stress. Graph of fitness values calculated by the rate of population change for each rDNA copy number test strain against the GFP competitor strain (180 rDNA copies) at 37° C (A) or grown in glycerol (B). Dots indicate individual fitness calculated of three (A) or two (B) replicates for each strain; shared dot and line colors indicate the experimental replicates.

To determine whether shifts in fitness among the rDNA copy number variants were specific to heat stress or common across other non-standard growth conditions, we performed competition experiments at 30°C in the nonfermentable carbon source glycerol. Results from these competitions differed from those obtained in standard laboratory condition and in response to heat stress (**Figure 4B**). The strain with the fewest rDNA copies (35) showed fitness comparable to the strain with 55 copies; strains carrying 80 and 100 copies performed only slightly better. We observed a fitness threshold at 140 copies, with the plateau of high fitness extending to strains with the longer rDNA arrays of 160 to 200 copies. We confirmed that rDNA copy numbers did not change during the competitions (**Figure S4**).

Spot assays examining the response of the rDNA copy number variants to heat stress, osmotic stress and ethanol found no differences among the tested strains; there was also no difference in their ability to acquire thermotolerance (**Figure S2**). Nevertheless, the results of our competition experiments strongly argue that rDNA copy number variation contributes to a strain’s fitness in response to different environments. Yeast cells face variable environments throughout their culturing in the laboratory (exponential growth, stationary phase, different carbon sources, freeze-thaw, etc.) and outside of it. Thus, the excess of rDNA copies not required for ribosome biogenesis in standard growth conditions may buffer condition-specific requirements for ribosome biogenesis and/or genome replication, the two processes previously identified as affected by rDNA copy number variation.

### A strain with substantially reduced rDNA copy number generates fewer petites than a control strain

Intrigued by the profound shift in fitness in response to glycerol, a nonfermentable carbon source, we examined mitochondrial function in the 35 rDNA copy number strain compared to the control 180 rDNA copy number strain. Specifically, we assessed mitochondrial function and mitochondrial genome stability by measuring the frequency of *petite* formation (Dimitrov *et al*. 2009). *Petites* are yeast colonies that have lost mitochondrial respiration function, and therefore are much smaller than respiration-competent, *grande* colonies on specialized glycerol media (**Figure 5A**). We performed *petite* frequency assays by selecting 15 medium-sized colonies from both strain backgrounds (35 rDNA copies, 180 rDNA copies), diluting them and plating approximately 350 cells onto glycerol-containing plates. After incubation for 5 days at 30°C, we counted all *petite* and *grande* colonies to determine the *petite* percentage for both strains.

**Figure 5:**
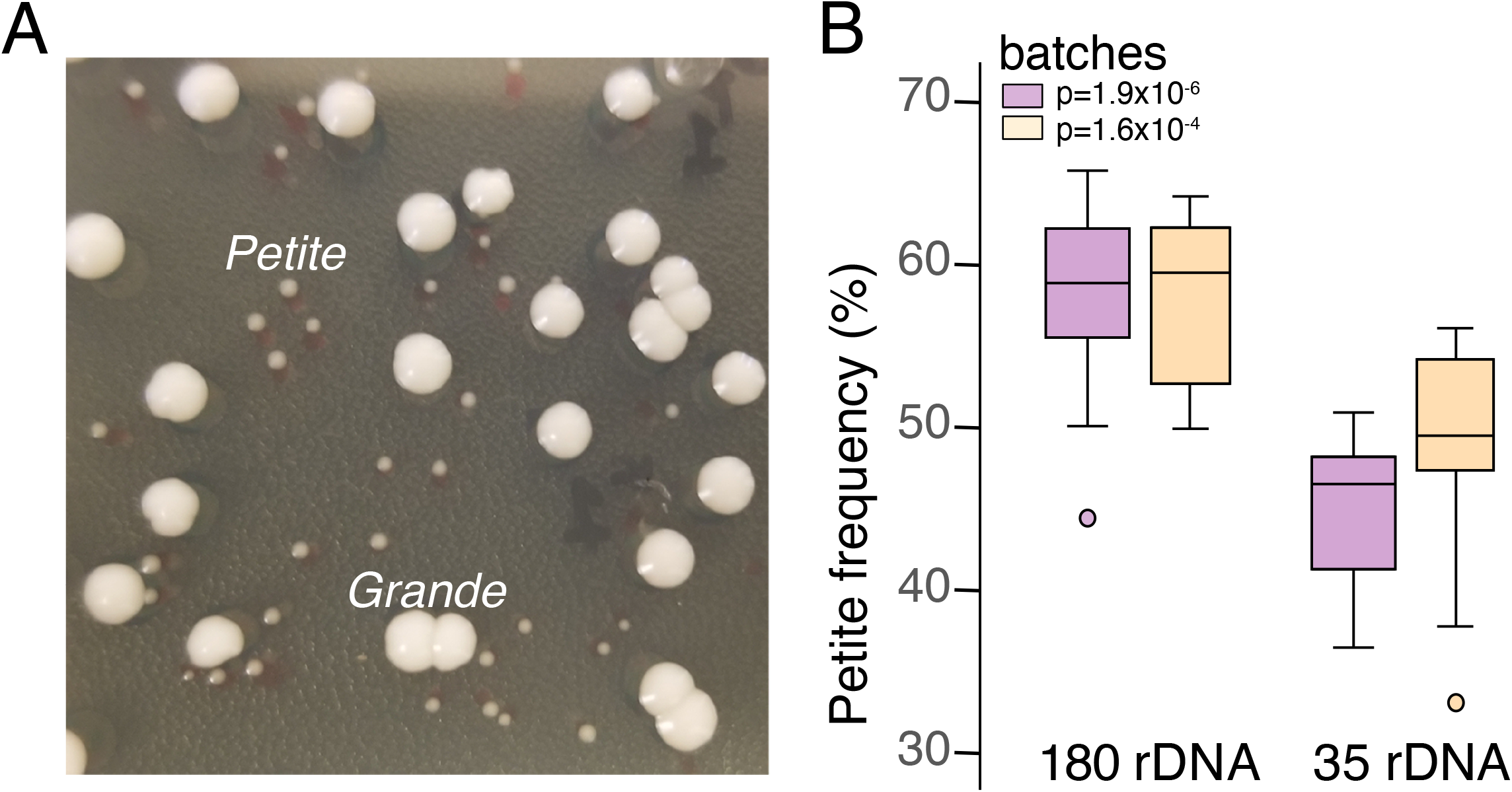
A strain with substantially reduced rDNA copy number generates fewer *petites* than a wild-type strain. (A) Approximately 350 cells from a single colony were plated onto YEPDG plates (0.1% glucose and 3% glycerol) to enhance *petite* vs. *grande* colony phenotypes (Dimitrov *et al*. 2009). For each strain, 15 plates were scored for total number of colonies and percentage of *petite* colonies. (B) The strain with 35 rDNA copies generated significantly fewer *petite* colonies than the wild-type strain in two replicate experiments.

The 35 rDNA copy number strain unexpectedly produced significantly fewer *petite* colonies (48.2% and 43.1% in the two replicates) than the 180 rDNA copy number strain (57.5% and 57.8%, respectively) (**Figure 5B**). Although this result appears inconsistent with the 35 rDNA copy number strain showing lower fitness than a 180 rDNA copy number variant in the glycerol competition experiments, it is possible that rDNA copy number variation affects these two traits in different ways.

## DISCUSSION

Here, we generated a set of rDNA copy number variants in otherwise isogenic *S. cerevisiae* strains to study the fitness consequences of rDNA copy number variation below and within the naturally occurring range of variation. In standard laboratory conditions, strains with the highest fitness had 98 to 160 rDNA copies, a range that overlaps with the distribution found in wild yeast isolates (Morton *et al*. 2020; Hall *et al*. 2022).

Strains with rDNA copy numbers either higher or lower than that range showed reduced fitness, with extreme variants, especially those with the lowest copy numbers, exhibiting the most severe fitness defects. However, rDNA copy number-dependent fitness was not static: the fitness of rDNA copy variants shifted in response to growth at increased temperature or in glycerol. We conclude that rDNA copy number variation plays a causal role in the response of yeast cells to different environments and that this variation should be considered when comparing strain responses across experimental growth conditions.

Upon re-introduction of *FOB1*, strains with 35 rDNA copies expanded their rDNA arrays to ~100 rDNA copies but further expansion stalled, consistent with a lack of fitness benefits with further expansion. Of the haploid wild yeast strains previously examined, 75% have rDNA copy numbers within the 98-160 fitness plateau (Morton *et al*. 2020; Hall *et al*. 2022). The remaining 25% of strains with higher rDNA copy numbers may experience more diverse environmental conditions than the other strains or carry genetic variants associated with rDNA copy expansion. Indeed, previous studies hint at a connection between rDNA copy number variation and genetic background. Kobayashi *et al*. report that a short rDNA strain fully expands its array to 150 copies after 150 generations (Kobayashi *et al*. 1998), far more rDNA copies than the ~100 we observed after 300 generations. The two studies differ in strain background: the strains in Kobayashi *et al*. 1998 were derived from W303 (Yano and Nomura 1991; Nogi *et al*. 1991) while our strains were derived from S288c. Although they are closely related laboratory strains, W303 and S288c differ in ~7000 non-synonymous polymorphisms (Liti *et al*. 2009; Matheson *et al*. 2017; Peter *et al*. 2018) and strains in the W303 background show higher rDNA copy numbers (250-300 copies) than those derived from S288c (140-150 copies) (Michel *et al*. 2005; Kwan *et al*. 2016; Lynch *et al*. 2019; Morton *et al*. 2020). The W303 strain might harbor genetic variants that facilitate rDNA expansion to higher copy numbers than in S288c strains, such as non-synonymous *SIR2* variants (Jack *et al*. 2015). Identifying these variants should further understanding of the regulation of rDNA copy number across all *S. cerevisiae* strains, and possibly in metazoans.

We show that environmental stress alters the relationship between rDNA copy number and fitness. The fitness plateau shifted to include higher rDNA copy numbers, and in glycerol, it excluded strains with 98 to 100 copies that showed high fitness in standard growth conditions. Second, the gradual fitness increase up to 98 copies found in standard laboratory conditions gave way to a markedly worse performance of rDNA copy number variants below the naturally occurring range. Thus, loss of rDNA copies, even by only a few copies, produced particularly strong deleterious effects in adverse conditions, and copy number at the higher end of the naturally occurring copy number range appear to buffer yeast cells against fluctuations in environment.

This interpretation is consistent with our observation that the S288c background slows rDNA array expansion at ~100 copies and shows high fitness at this copy number, yet the S288c strain typically contains ~150 rDNA copies (Ide *et al*. 2013; Kwan *et al*. 2016; Morton *et al*. 2023). Since the fitness plateau shifted to higher copy numbers in response to stress, the strain may have been selected under laboratory conditions for 150 rDNA copies rather than ~100 copies through exposure to a myriad of intermittent stresses: *e*.*g*., cold/freezing temperatures, heat shock, reduced nutrient availability (Kwan *et al*. 2016). We speculate that conducting the expansion experiment in fluctuating environments might have resulted in rDNA arrays of ~150 copies or higher; however, it is non-trivial to design such an experiment to be informative and artifact-free.

At higher rDNA copy numbers, such as 180 and 200, fitness can decrease in standard growth conditions, which may reflect the more sparse representation of wild yeast isolates with higher rDNA copy numbers. The fitness reduction in the 180 and 200 rDNA copy number strains seems inconsistent with a previous model that implicated the excess of rDNA copies over the number required for ribosome biogenesis in maintaining rDNA and genome stability (Ide *et al*. 2010). If this were the case, the additional rDNA copies would be expected to increase fitness in standard growth conditions rather than reduce it. A possible mechanism explaining the reduced fitness in strains with increased rDNA copy number is the cost of maintaining the additional copies, especially since an extra 25 rDNA copies (~227 kb) represent a similar amount of DNA as the smaller *S. cerevisiae* chromosomes I (230 kb) and III (316 kb) (Cherry *et al*. 2012). Dedicating resources to maintaining an additional chromosome’s worth of repetitive, presumably silenced, DNA could plausibly reduce fitness (Sunshine *et al*. 2015). Further studies with strains containing a wider range of rDNA copy numbers above the natural range are needed to distinguish among possible fitness effects, although such strains are not easy to construct.

The phenotypic consequences of rDNA copy number variation has been increasingly studied in multicellular animals and plants (Xu *et al*. 2017; Kasselimi *et al*. 2022; Hall *et al*. 2022; Morton *et al*. 2023). In the nematode *C. elegans*, individual animals with rDNA copy numbers below the naturally occurring range display a gradient of developmental defects, ranging from subtle developmental delays to developmental arrest and strikingly variable morphological defects in postembryonic development (Cenik *et al*. 2019; Morton *et al*. 2023). These observations suggest that specific stages of development and specific tissues have their own rDNA copy number requirements, perhaps because each stage and tissue represent different cellular environments. That rDNA copy number-dependent fitness shifts in yeast with environmental conditions may be analogous; although yeast cells do not form specific tissues, they experience a wide range of environmental conditions.

The shifts in environmental response of cells with different rDNA copy number are particularly intriguing in the context of cancer. Reduction of rDNA copy number has been reported in mTOR-related tumors compared to healthy, non-cancerous tissue (Xu *et al*. 2017; Kasselimi *et al*. 2022). Because rDNA copy number reduction causes defects in genome replication and cell cycle control in yeast (Ide *et al*. 2010; Kwan *et al*. 2023), rDNA copy number reduction in human cells may contribute to the characteristic loss of genome stability that precedes cancer pathogenesis (Hanahan and Weinberg 2011). Moreover, the reduction in rDNA copy number might shift the fitness of pre-cancerous and/or cancer cells in response to the stresses of host environment or treatment, analogous to our observations in yeast.

## Data availability

Transcriptome raw sequencing reads can be found at the NCBI short read archive under the bioproject PRJNA1061514.

## Acknowledgements

This work was supported by the following funding sources: NIGMS R01 GM122088 and R35 GM139532 and NHGRI grant RM1 HG010461to CQ, and NIGMS R35 GM122497 to BJB. Flow cytometry reported in this work was performed at the DLMP Flow Cytometry Core at the University of Washington.

## FIGURE LEGENDS

**Figure S1: Fitness trends from rDNA copy number variation are highly reproducible.**

(A) Growth rates for subset of rDNA copy number variant strains used for fitness competition experiments. Doubling time is indicated in parentheses. No significant difference is growth rate was detected in standard growth conditions. (B) CHEF gel electrophoresis and (C) Southern blotting using an rDNA probe (*NTS2*) from two additional batches presented in Figure 1. The two chromosome XII bands from both rDNA copy number variant test strain and GFP competitor are both visible in the last day competition culture. No change to rDNA copy number was detected.

**Figure S2: Examination of rDNA copy number variants by spot assays for response to stress.**

No difference was detected between rDNA copy number variants by spot assays examining (A) thermotolerance on both normal YEPD and glycerol media or (B) salt stress and ethanol stress. (C) No difference in acquisition of thermotolerance was observed by spot assays. Acquisition of thermotolerance was examined by survival of strains pre-treated with a mild heat shock at 37ºC for 60 minutes. Both pre-treated and untreated cells were then exposed to extreme heat shock (50ºC) for 4, 8, and 12 minutes before spotting onto plates. Plates were grown at 30ºC for two days.

**Figure S3: Heat stress fitness trends due rDNA copy number variation are reproducible.**

(A) CHEF gel and (B) Southern blot (*NTS2*) of competition culture sample on the last day for both replicates from 37° C competition experiments. No change in rDNA copy number was found. (E) Growth rates for subset of rDNA copy number variant strains used for fitness competition experiments. Doubling time is indicated in parentheses. No significant difference in growth rate was detected at 37ºC.

**Figure S4: Glycerol rDNA copy number.**

(A, C) CHEF gel and (B, D) Southern blot (*NTS2*) of competition culture sample on the last day for both replicates grown in glycerol. No change in rDNA copy number is detected.

## METHODS

### Yeast strains

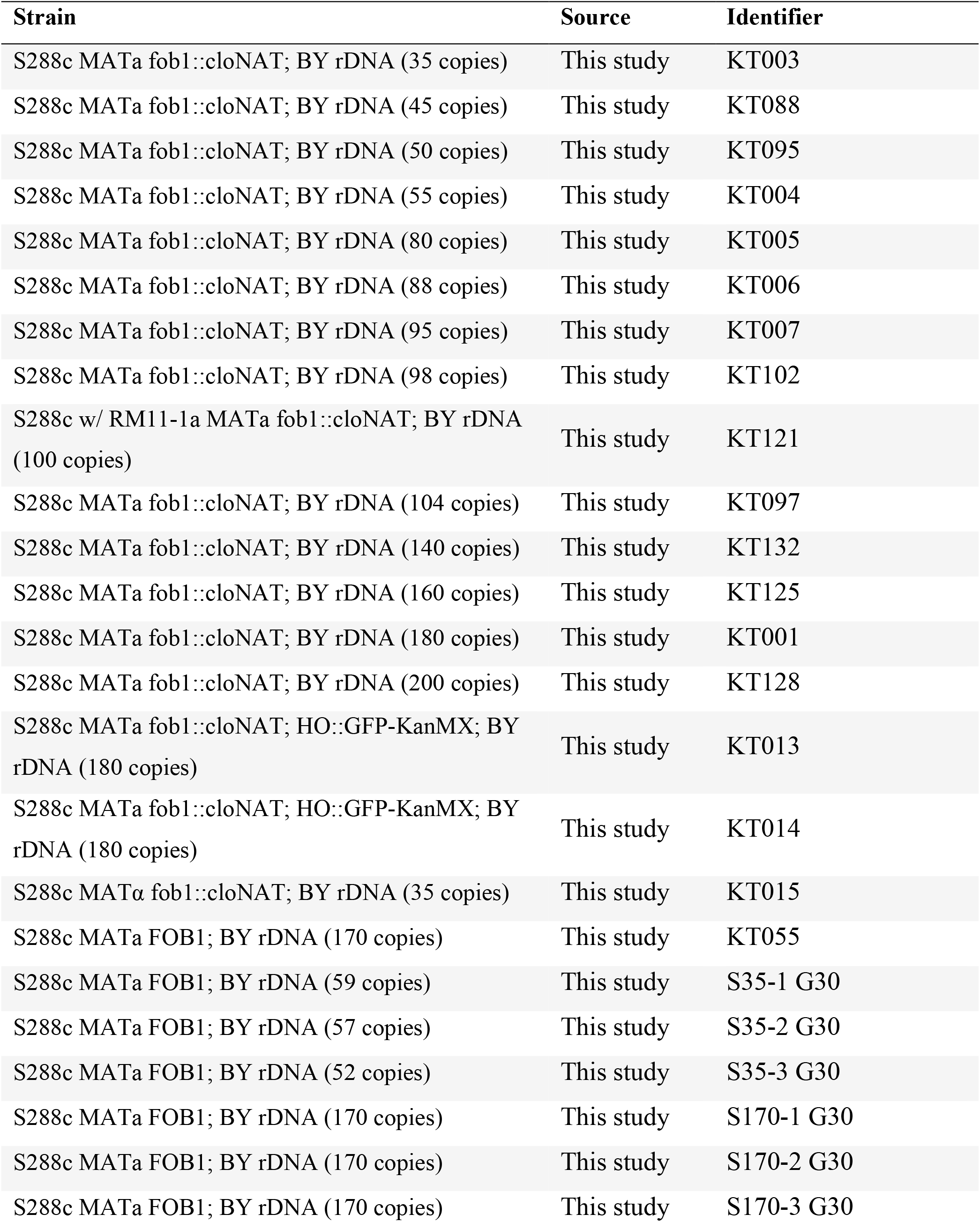

### Yeast media

Yeast strains used in 30°C and 37°C yeast competition assays were grown in synthetic complete media buffered with 1% succinic acid (per liter: 1.45 g yeast nitrogen base, 20 g glucose, 10 g succinic acid, 6 g NaOH, 5 g (NH4)_2_SO_4_, 2.8 g amino acid powder mix with pH adjusted to 5.8).

Yeast strains used in glycerol yeast competition assays were grown in synthetic complete media buffered with 1% succinic acid (per liter: 1.45 g yeast nitrogen base, 30 g glycerol, 10 g succinic acid, 6 g NaOH, 5 g (NH4)_2_SO_4_, 2.8 g amino acid powder mix with pH adjusted to 5.8).

Yeast strains used in caloric restriction yeast competition assays were grown in synthetic minimum media buffered with 1% succinic acid (per liter: 1.61 g yeast nitrogen base, 1 g glucose, 11.10 g succinic acid, 6.67 g NaOH, 100 mg (NH4)_2_SO_4_ with pH adjusted to 5.8).

### Preparation of DNA embedded in agarose

Cells were inoculated into 2 mL synthetic complete media buffered with 1% succinic acid and allowed to grow overnight to stationary phase (1.5 – 3 × 10^8^ cells/mL). 200 μL of cells were pelleted for 1 minute at 15,000 rcf and the supernatant was discarded. Cells were washed with 85 μL 50 mM EDTA, resuspended in 90 μL 1% SeaPlaque™ GTG™ agarose in 50 mM EDTA then transferred into plug molds. Plugs were allowed to solidify for 15 minutes at 4ºC then incubated in 1 mL spheroplasting solution (1.0 M sorbitol, 20 mM EDTA pH 8.0, 10 mM Tris-HCl pH 7.4, 14.3 mM β-mercaptoethanol, 0.5 mg/mL Zymolyase-20T [Amsbio]) for 2 – 4 hours at 37ºC with gentle shaking. Plugs were washed once with LDS (1 % lithium dodecyl sulfate, 100 mM EDTA pH 8.0, 10 mM Tris–HCl pH 8.0) and incubated overnight at 37ºC in LDS. Plugs were then washed 3 × 20 minutes in 0.2X NDS (1X NDS pH 9.5: 0.5 M EDTA, 10 mM Tris base, 1% Sarkosyl) and 5 × 20 minutes in TE pH 8.0. All processed plugs were stored at 4ºC in TE pH 8.0 until use.

### CHEF gel analysis

Intact chromosomes were resolved by contour-clamped homogeneous electric field (CHEF) gel electrophoresis. A small slice (5 mm x 2 mm x 3 mm) of all genomic DNA agarose plugs were embedded in a 0.8% low electroendosmosis (LE) agarose gel containing filtered 0.5X TBE. CHEF gels were run in 2.3 L of 0.5X TBE using a Bio-Rad CHEF-DRII electrophoresis cell at 100V for 68 hours (switch time = 300 to 900 seconds). All gels were stained with ethidium bromide to visualize chromosome XII, which contains the rDNA, and all other chromosomes. *Hansenula wingei* (*H. wingei*) chromosomal DNA size marker standards were included in each CHEF gel electrophoresis run for size comparison.

### Southern blotting

All CHEF gels were transferred to Genescreen Hybridization membrane using standard Southern blotting protocols (Tsuchiyama et al., 2013). We then hybridized the sequences of interest using a ^32^P-labeled probe. The blots were exposed to X-ray film and to Bio-Rad Molecular Imaging FX phosphor screens for visualization and quantification of signal intensity.

### Yeast competitions

Cells were streaked out on YEPD plates and allowed to incubate overnight at 30°C. Cells were inoculated into 2 mL synthetic complete media with 2% glucose buffered with 1% succinic acid and allowed to grow overnight to stationary phase (1.5–3 × 10^8^ cells/mL). 100 μL of overnight cultures was transferred to 5 mL synthetic complete media (1:50 dilution) and allowed to grow at 30°C for ~5 hours. After the 5 hour incubation period, 500 μL of test strain cultures were mixed with 500 μL of 180 rDNA GFP competitor strain culture and vortexed thoroughly. 500 μL of each test strain and GFP competitor strain culture mix was transferred to 1.5 mL sterile water (1:4 dilution) to record cell number of all strains with the Gilford Stasar Spectrophotometer at a wavelength 660. 5 μL of each test strain and GFP competitor strain culture mix was transferred to 5 mL synthetic complete media (1:1000 dilution) and incubated at 30°C overnight. Cell fitness was determined by measuring the change in GFP population of cells over time with flow cytometry of cell culture aliquots from day zero and day five. All fitness defects and advantages seen in strains used in these experiments were reproducible across several competition experiments.

### Flow cytometry

~350 μL of day zero yeast competition experiment cultures were suspended to 1 mL 50 mM sodium citrate and sonicated in preparation of flow cytometry. 100 μL of day one through day five yeast competition experiment overnight cultures were suspended in 1 mL 50 mM sodium citrate and sonicated in preparation of flow cytometry. Cells were analyzed on a BD Canto II flow cytometer and flow cytometry data was analyzed using FlowJo software.

### RNA extraction for RNA-seq

Asynchronous and late S phase logarithmic phase cells were collected, and genomic DNA was isolated using an acid phenol: chloroform extraction protocol by resuspending frozen pellets in 200 μL lysis buffer (10 mM Tris, pH 8.0, 10 mM EDTA, 5% SDS), and 200 μL acid phenol and vortexed for 2 min. Cells were resuspended and incubated at 65°C for 1 hour and vortexed occasionally. Incubated samples were kept on ice for 10 minutes before centrifuging samples at 4°C at 15,000 rcf for 10 minutes. After transferring the aqueous layer to new 1.5 microcentrifuge tubes, we added equal volumes of chloroform, and vortexed vigorously before centrifuging samples at 15,000 rcf for 10 minutes. To remove bulk of ribosomal RNA (rRNA) before sequencing library prep, DNA oligos complimentary to rRNA sequences were incubated with RNA samples and then treated with RNase H to degrade RNA:DNA hybrids. RNA was then purified again using phenol:chloroform and ethanol precipitated. rRNA-depleted RNA was purified with a 2.2X RNAClean SPRI bead treatment (Beckman Coulter). Purified, rRNA-depleted RNA was treated with Turbo DNase (Invitrogen) to remove the rRNA-hybridizing DNA oligos: 0.1 volumes (2 μL) of 10X TURBO DNase Buffer was added to the RNA, followed by 2 μL TURBO DNase. The mixture was incubated at 37°C for 45 minutes. The DNase-treated RNA was then cleaned with 2.2X RNA SPRI beads and a total volume of 12 μL was eluted. 9 μL of the eluted volume was used immediately for poly-A mRNA capture.

9 μL of final DNase-treated RNA and 16 μL nuclease-free ultrapure water were added to 25 μL RNA Purification Beads, mixed and incubated at 65°C for 5 minutes then at 4°C for 30 seconds and finally at 23°C for 5 minutes. Samples were placed on a magnetic stand, supernatant was discarded, and 100 μL Bead Washing Buffer was added. After removing all residual supernatant, 25 μL Elution Buffer was added to samples and were incubated at 80°C for 2 minutes. 25 μL Bead Binding Buffer to samples and then washed with 100 μL Bead Washing Buffer. After removing supernatant, 19 μL ice cold Fragmentation Master Mix (10.5 μL Nuclease-free ultrapure water and 10.5 μL Elute, Prime, Fragment High Mix per sample) was added to each sample and incubated at 94°C for 8 minutes. 17 μL of mRNA PCR product was added to new PCR tubes on ice in preparation of First strand cDNA synthesis.

First strand cDNA synthesis was performed by adding 8 μL First Strand Synthesis Master (9 μL First Strand Synthesis Act D Mix and 1 μL Reverse Transcriptase per sample) to 17 μL of each mRNA PCR product and incubated at 25°C for 10 minutes then at 42°C for 15 minutes and finally at 70°C for 15 minutes. Second strand cDNA synthesis was performed by adding 25 μL of the first cDNA strand reaction products with 25 μL Second Strand Marking Master Mix and incubated at 16°C for 1 hour. After the incubation period, second cDNA strand reaction products were captured with 90 μL AMPure XP beads, supernatant was discarded, and AMPure XP beads were cleaned up twice with 175 μL fresh 80% ethanol on a magnetic stand. After removing all residual ethanol, AMPure XP beads were resuspended in 19.5 μL Resuspension Buffer and 17.5 μL supernatant was transferred into new PCR tubes. The first and second cDNA products were used immediately or stored at −20°C overnight for use the next day.

In preparation for adenylating 3’ ends, 12.5 μL A-Tailing Mix was added to the first and second cDNA products and incubated at 37°C for 30 minutes then at 70°C for 5 minutes. Anchors were ligated to samples by adding 2.5 μL Resuspension Buffer, 2.5 μL RNA Index Anchors (Illumina), 2.5 μL Ligation Mix and incubated at 30°C for 10 minutes followed by adding 5 μL Stop Ligation Buffer. 34 μL AMPure XP beads were added to samples, supernatant was discarded, and AMPure XP beads were cleaned up twice with 175 μL fresh 80% ethanol on a magnetic stand. After removing all residual ethanol, AMPure XP beads were resuspended in 22 μL

Resuspension Buffer and 20 μL supernatant was transferred into new PCR tubes. The ligated cDNA products were used immediately or stored at −20°C overnight for use the next day.

Dual-indexed libraries were generated by adding 10 μL index adapters (Illumina) and 20 μL Enhanced PCR Mix to the ligated cDNA products. The mixture was mixed and incubated for 13 cycles at 98°C for 10 seconds, 60°C for 30 seconds, and 72°C for 30 seconds followed by an incubation at 72°C for 5 minutes. 50 μL AMPure XP beads were added to samples, supernatant was discarded, and AMPure XP beads were cleaned up twice with 175 μL fresh 80% ethanol on a magnetic stand. After removing all residual ethanol, AMPure XP beads were resuspended in 17 μL Resuspension Buffer and 15 μL supernatant was transferred into new PCR tubes. The concentrations and quality of all libraries were examined for quality using TapeStation 2200 (Agilent Technologies) and Qubit (Thermo Fisher Scientific) then subsequently diluted to the appropriate starting concentrations for sequencing.

Libraries were sequenced using a NextSeq 550 with a 75 Hi kit (Illumina). Read lengths used were: Index 1: 8bp, Index 2: 8bp, Read 1: 38bp, Read 2: 38bp.

### Transcriptome profiling

We aligned to the S288C reference genome version R64.1.1 and obtained gene counts using hisat2 version 2.2.1 (Kim et al., 2019). DESeq2 was used to identify differentially expressed genes (Love et al., 2014). Genes with an adjusted p-value of less than 0.01 were further analyzed: six that differed between the 35- and 180-rDNA copy number in asynchronous culture (*PRP11, HSP12, STR3, NCE103, YGP1*, and *CIN5*) and 708 that differed between the 35- and 180-rDNA copy number in synchronous culture during late S phase, with three genes in common between these two sets (*PRP11, STR3*, and *HSP12*).

We performed gene ontology enrichment analysis for the following four sets of genes, each of which consists of only genes determined to be differentially expressed in late S phase between the 35- and 180-rDNA copy number strains: (1) genes that were more expressed in the 35-rDNA copy number strain than the 180-rDNA copy number strain; (2) genes that were more expressed in the 180-rDNA copy number strain than the 35-rDNA copy number strain; (3) genes that were at least 1.5x more expressed in the 35-rDNA copy number strain than the 180-rDNA copy number strain; (4) genes that were at least 1.5x more expressed in the 180-rDNA copy number strain than the 35-rDNA copy number strain. Set (1) was enriched for ribosomal components, for example 110 of the 420 genes in this set were in the “cytosolic ribosome” category (GO:0022626; enrichment padj<1e-95). Set (2) was enriched for proteolysis-related genes, for example 60 of the 288 genes in this set were in the “proteolysis involved in protein catabolic process” category (GO:0051603; enrichment padj<1e-22). Set (3) was enriched for chromatin-related genes, for example 8 of the 88 genes in this set were in the “structural constituent of chromatin” category (GO:0030527; enrichment padj<1e-10). Set (4) was enriched for stress-related genes, for example 36 of the 83 genes in this set were in the “response to stress” category (GO:0006950; enrichment padj<1e-7). We colored each differentially expressed gene that was a member of any of these four enriched GO categories, regardless of whether the gene is in the set enriched for that GO category.

### Spot assays

Cells were grown to log-phase, diluted in sterile water in 3-fold dilutions starting with a cell concentration of 4 × 10^5^ cells/mL. 2.5 μL was spotted onto YEPD (1% yeast extract, 2% peptone, and 2% glucose), YEPD + 0.7 M Sodium chloride, YEPD + 3% ethanol, YEPD + 6% ethanol, and YEPG plates. All plates were scanned after 48 hours of growth at 30°C. Thermotolerance assays were adapted from Lindquist & Kim 1996.

### Petite frequency assays

The procedure adapted from Dimitrov *et al*. 2009 was used. Medium sized colonies from on direct-from-freezer-stock streakouts on YEPD plates were inoculated 2 mL cultures tubes filled with WF-N media. A portion of 2 mL culture was used to determine cell densities of cultures and diluted to 1 × 10^5^ cells/mL. Cultures were diluted again to a cell concertation of 2.5 × 10^3^ cells/mL. Diluted cultures were sonicated and 150 μL was plated onto YEPDG (1% yeast extract, 2% peptone, 0.1% glucose, and 3% glycerol) plates and allowed to grow at 30°C for 5 days. After the incubation period, all plates were scored for *petite* and *grande* colonies.

